# Unique Peptide Signatures Of SARS-CoV-2 Against Human Proteome Reveal Variants’ Immune Escape And Infectiveness

**DOI:** 10.1101/2021.10.03.462911

**Authors:** Vasileios Pierros, Evangelos Kontopodis, Dimitrios J. Stravopodis, George Th. Tsangaris

**Affiliations:** Proteomics Research Unit, Biomedical Research Foundation of the Academy of Athens; 11527 Athens, Greece; Section of Cell Biology and Biophysics, Department of Biology, School of Science, National and Kapodistrian University of Athens; 15701 Athens, Greece

**Keywords:** SARS-CoV-2, COVID-19, Core Unique Peptides, Uniquome, Spike protein, NF9 peptide, Mutations, Delta variant, Immune escape, Infectiveness

## Abstract

SARS-CoV-2 pandemic has emerged the necessity of the identification of sequences sites in viral proteome appropriate as antigenic sites and treatment targets. In the present study, we apply a novel approach for deciphering the virus-host organism interaction, by analyzing the Unique Peptides of the virus with a minimum amino acid sequence length defined as Core Unique Peptides (CrUPs) not of the virus *per se*, but against the entire proteome of the host organism. The result of this approach is the identification of the CrUPs of the virus itself, which do not appear in the host organism proteome. Thereby, we analyzed the SARS-CoV-2 proteome for identification of CrUPs against the Human Proteome and they are defined as C/H-CrUPs. We found that SARS-CoV-2 include 7.503 C/H-CrUPs, with the SPIKE_SARS2 being the protein with the highest density of C/H-CrUPs. Extensive analysis indicated that the P681R mutation produces new C/H-CrUPs around the R685 cleavage site, while the L452R mutation induces the loss of antigenicity of the NF9 peptide and the strong(er) binding of the virus to ACE2 receptor protein. The simultaneous existence of these mutations in variants like Delta results in the immune escape of the virus, its massive entrance into the host cell, a notable increase in virus formation, and its massive release and thus elevated infectivity.

## Main Text

Covid-19 pandemic has emerged the urgent necessity of the identification of sequence sites of the SARS-CoV-2 viral proteome that serve as appropriate treatment targets and antigenic sites suitable for production of therapeutic vaccines.

In our previous studies, we have defined as Unique Peptides (UPs) the peptides that their amino acid sequence appears only in one protein across a given proteome. We have also introduced the term of Core Unique Peptides (CrUPs), which are the peptides with a minimum amino acid sequence length that appear only in one protein across a given proteome, being thus a unique signature for the particular protein identification (Alexandridou et al., 2009; Kontopodis et al., 2019). Thereby, to map the UP landscape of a proteome under examination, we have herein developed a novel and advanced bioinformatics tool including big data analysis (Supp. Materials and Methods). Its employment to analysis of the 20.430 reviewed *Homo sapiens* proteins resulted in the identification of 7.263.888 CrUPs, which are parts of the Human Uniquome that is defined as the total set of unique peptides belonging to the Human proteome (Kontopodis et al., 2019).

Recently, in order to elucidate the virus-host organism interaction, we designed an advanced bioinformatics approach to analyze the CrUPs of the virus against the host organism proteome. These peptides are different from the virus CrUPs *per se*, which are defined as the minimum amino acid sequence length peptides appeared only in one protein across the virus proteome. The virus CrUPs against the host organism proteome have two distinct properties: (a) they are unique in the virus proteome and (b) they do not exist in the host organism proteome. Based on these properties, the virus CrUPs against the host organism proteome illuminate our knowledge about the virus-host interaction, the infectiveness and the pathogenicity of the virus, and, most importantly, they can be used as antigenic and diagnostic peptides, and possible treatment targets. Furthermore, these unique peptides constitute a completely new entity of peptides able to advance our knowledge about the construction of viral and Human Uniquomes (Kontopodis et al., 2019).

Since human cells can host the SARS-CoV-2 virus, we have herein engaged our novel bioinformatics platform not only for the profiling of CrUPs in SARS-CoV-2 proteome *per se*, but, most importantly, for their identification against the human proteome (C/H-CrUPs). Remarkably, C/H-CrUPs can likely serve as targets for the immune response upon infection, and antigenic sites with major pharmaceutical and diagnostic potential for the successful clinical management of the Covid-19 pandemic.

The SARS-CoV-2 proteome is structurally quite simple. In the UNIPROT database (version 7/2021), 16 reviewed and 75.714 unreviewed proteins have been included (Jungreis et al., 2021). For the present study, only the 16 reviewed proteins are examined, since the unreviewed proteome components contain (among others) duplicate registrations, unverified sequences and protein fragments, which could lead to unreliable data regarding the uniqueness of a protein sequence.

To recognize all the CrUPs being embraced in SARS-CoV-2 proteome against the human proteome, we *in silico* constructed a new, artificial, “hybrid-proteome” that contained all the reviewed human proteins (20.430 proteins) plus the one protein derived from SARS-CoV-2 viral proteome (20.431 proteins). Thus, 16 “hybrid proteomes” including the 16 SARS-CoV-2 proteins were constructed. Hence, these “hybrid proteomes” were bioinformatically searched one by one for the identification of SARS-CoV-2-specific CrUPs in human protein sequence environments (C/H-CrUPs).

Strikingly, 7.503 C/H-CrUPs were detected, with 4.213 of them being presented one time in the SARS-CoV-2 proteome, 3.289 being observed two times in the viral proteome and only one peptide (“VNNATN”) with a 6 amino acid length being recognized three times (Table 1 and Data S1). Data processing and analysis unveiled that C/H-CrUPs retain a length range from 4 to 9 amino acids, while longer peptides could not be identified in the SARS-CoV-2 virus proteome. Length distribution showed that the majority of C/H-CrUPs have a 6 amino acid length, whereas only one with 4 amino acids and only two with 9 amino acids C/H-CrUPs were observed (Fig 1).

**Table 1:**
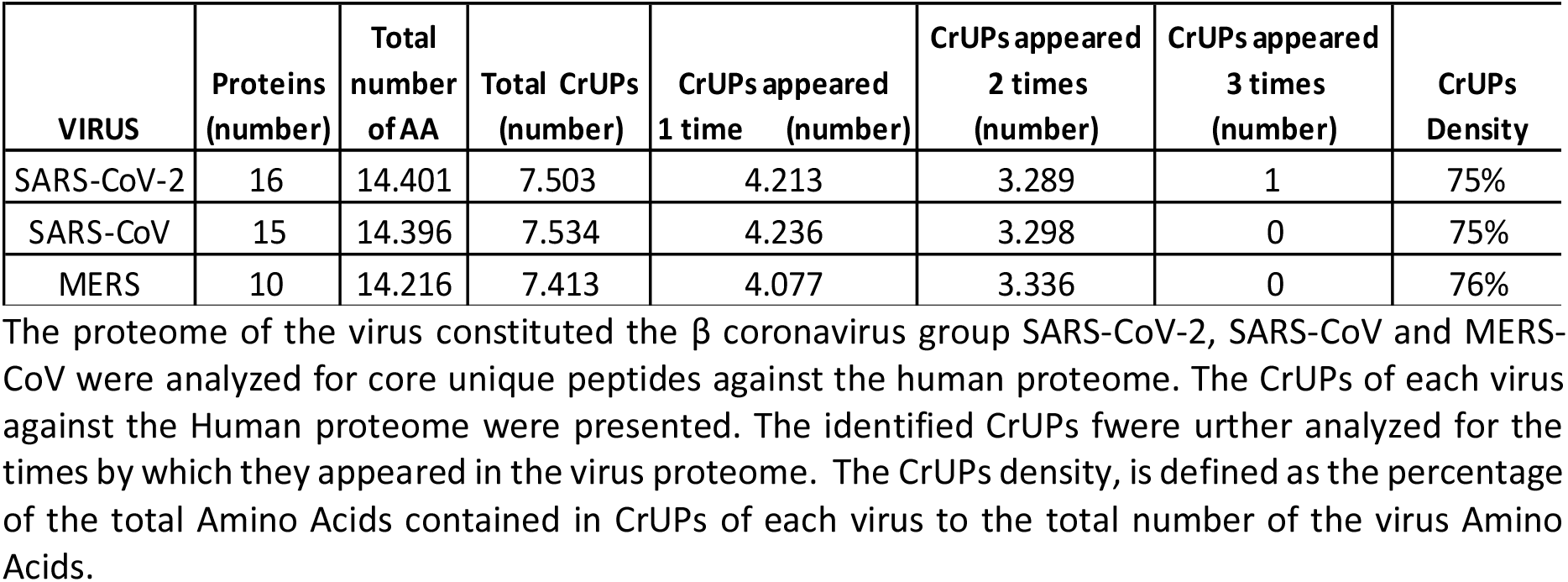
Viruses CrUPs against Human proteome.

| VIRUS | Proteins (number) | Total number of AA | Total CrUPs (number) | CrUPs appeared 1 time (number) | CrUPs appeared 2 times (number) | CrUPs appeared 3 times (number) | CrUPs Density |
| --- | --- | --- | --- | --- | --- | --- | --- |
| SARS-CoV-2 | 16 | 14.401 | 7.503 | 4.213 | 3.289 | 1 | 75% |
| SARS-CoV | 15 | 14.396 | 7.534 | 4.236 | 3.298 | 0 | 75% |
| MERS | 10 | 14.216 | 7.413 | 4.077 | 3.336 | 0 | 76% |

The proteome of the virus constituted the $\beta$ coronavirus group SARS-CoV-2, SARS-CoV and MERS-CoV were analyzed for core unique peptides against the human proteome. The CrUPs of each virus against the Human proteome were presented. The identified CrUPs were further analyzed for the times by which they appeared in the virus proteome. The CrUPs density, is defined as the percentage of the total Amino Acids contained in CrUPs of each virus to the total number of the virus Amino Acids.

**Figure 1.**
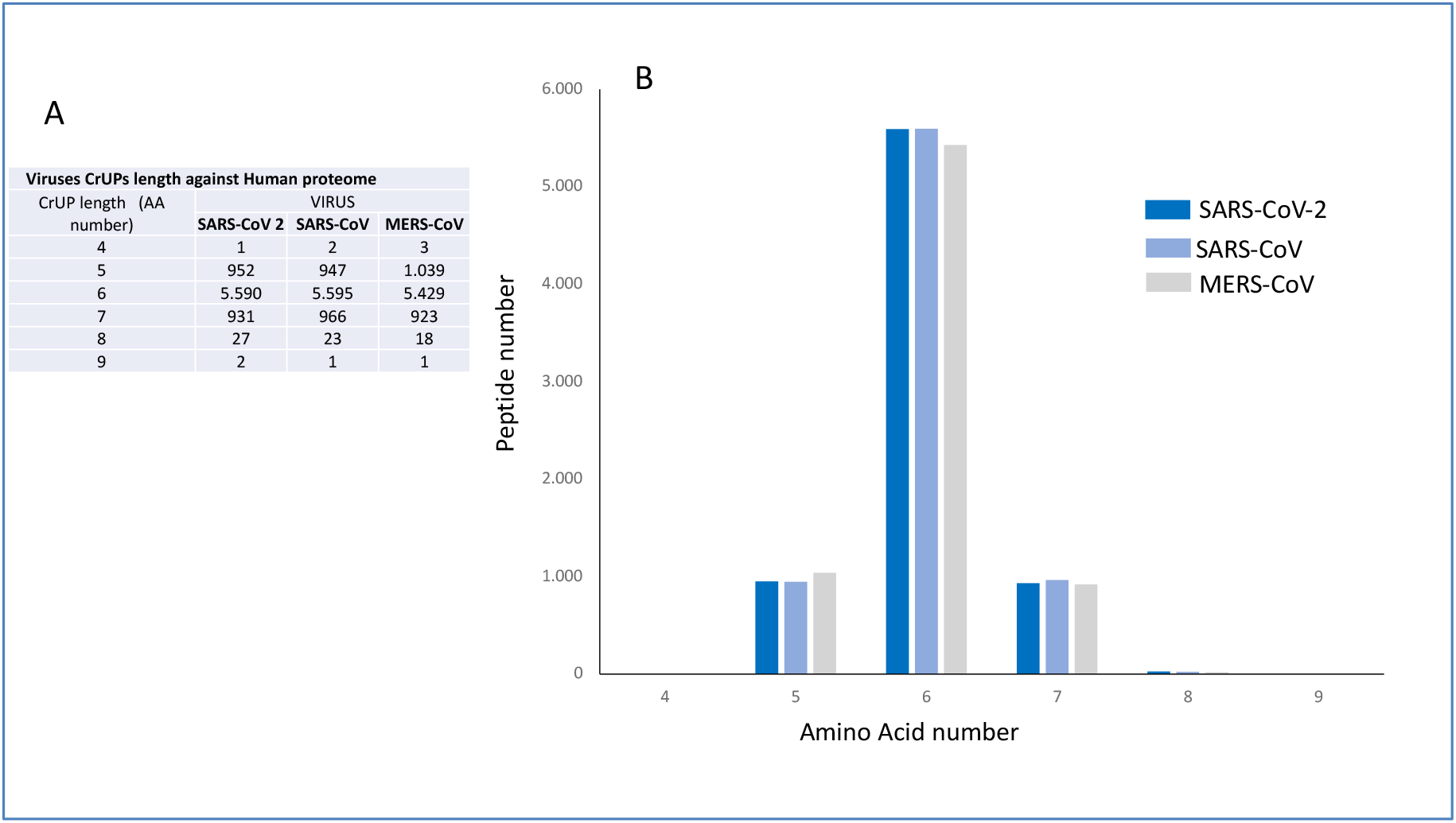
Amino acid length distribution of viruses Core Unique Peptides (CrUPs) against Human proteome. **A)** Table of the CrUPs of SARS-CoV-2, SARS-CoV and MERS-CoV viruses against the Human proteome. The CrUPs were identified, listed and grouped according to their amino acid length. **B)** Graphical representation of the CrUPs amino acid length across β coronavirus group.

The distribution of C/H-CrUPs across SARS-CoV-2 proteins demonstrated that the Replicase Polyprotein 1ab (R1AB_SARS2), which is the longest viral protein consisted of 7.096 amino acids, produces almost half of the identified C/H-CrUPs (5.334; 49,3%) (Table 2). On the other hand, the Putative ORF3b protein (ORF3B_SARS2), with a length of 22 amino acids, produces only 15 C/H-CrUPs that show a protein density of 68%. Notably, Spike glycoprotein (SPIKE_SARS2) is presented with the highest C/H-CrUPs density (78%), thus indicating its intriguing feature to carry the highest number of C/H-CrUPs (987), in terms of their physical length, as opposed to the ORF3c protein (ORF3C_SARS2), which is characterized by a respective density of only 56% (Table 2). A typical example for the construction of C/H-CrUPs is the peptide “PDEDEEEGD”. This peptide is a 9 amino acid in length C/H-CrUP that belongs to Replicase polyprotein 1a (R1A_SARS2), starting at position 927 and ending at position 935 (Fig S1). Around this peptide, 8 C/H-CrUPs were recognized with a 5-7 amino acid length range.

**Table 2.** Viruses detailed analysis.

| **SARS-CoV-2** |  |  |  |  |  |
| --- | --- | --- | --- | --- | --- |
| **Entry ID** | **Entry Name** | **Protein Name** | **Length**<br>**(AA number)** | **C/H-CrUPs**<br>**(number)** | **C/H-CrUPs**<br>**Density** |
| P0DTD1 | R1AB_SARS2 | Replicase polyprotein 1ab | 7096 | 5334 | 75% |
| P0DTC1 | R1A_SARS2 | Replicase polyprotein 1a | 4405 | 3294 | 75% |
| P0DTC2 | SPIKE_SARS2 | Spike glycoprotein | 1273 | 987 | **78%** |
| P0DTC9 | NCAP_SARS2 | Nucleoprotein | 419 | 308 | 74% |
| P0DTC3 | AP3A_SARS2 | ORF3a protein | 275 | 210 | 76% |
| P0DTC5 | VME1_SARS2 | Membrane protein | 222 | 171 | 77% |
| P0DTC7 | NS7A_SARS2 | ORF7a protein | 121 | 90 | 74% |
| P0DTC8 | NS8_SARS2 | ORF8 protein | 121 | 82 | 68% |
| P0DTD2 | ORF9B_SARS2 | ORF9b protein | 97 | 69 | 71% |
| P0DTD3 | ORF9C_SARS2 | Putative ORF9c protein | 73 | 50 | 68% |
| P0DTC4 | VEMP_SARS2 | Envelope small membrane protein | 75 | 48 | 64% |
| P0DTC6 | NS6_SARS2 | ORF6 protein | 61 | 44 | 72% |
| P0DTG0 | ORF3D_SARS2 | Putative ORF3d protein | 57 | 40 | 70% |
| P0DTD8 | NS7B_SARS2 | ORF7b protein | 43 | 29 | 67% |
| P0DTG1 | ORF3C_SARS2 | ORF3c protein | 41 | 23 | 56% |
| P0DTF1 | ORF3B_SARS2 | Putative ORF3b protein | 22 | 15 | 68% |
| **SARS-CoV** |  |  |  |  |  |
| **Entry ID** | **Entry Name** | **Protein Name** | **Length**<br>**(AA number)** | **S/H-CrUPs**<br>**(number)** | **S/H-CrUPs**<br>**Density** |
| P0C6X7 | R1AB_SARS | Replicase polyprotein 1ab | 7.073 | 5.346 | 76% |
| P0C6U8 | R1A_SARS | Replicase polyprotein 1a | 4.382 | 3.301 | 75% |
| P59594 | SPIKE_SARS | Spike glycoprotein | 1.275 | 970 | **76%** |
| P59595 | NCAP_SARS | Nucleoprotein | 422 | 319 | 76% |
| P59632 | AP3A_SARS | ORF3a protein | 274 | 208 | 76% |
| P59596 | VME1_SARS | Membrane protein | 221 | 162 | 73% |
| P59633 | NS3B_SARS | ORF3b protein | 154 | 113 | 73% |
| P59635 | NS7A_SARS | ORF7a protein | 122 | 93 | 76% |
| P59636 | ORF9B_SARS | ORF9b protein | 98 | 71 | 72% |
| Q80H93 | NS8B_SARS | ORF8b protein | 84 | 59 | 70% |
| P59637 | VEMP_SARS | Envelope small membrane protein | 75 | 47 | 63% |
| Q7TLC7 | Y14_SARS | Uncharacterized protein 14 | 70 | 45 | 64% |
| P59634 | NS6_SARS | ORF6 protein | 63 | 44 | 70% |
| Q7TFA1 | NS7B_SARS | Protein non-structural 7b | 44 | 27 | 61% |
| Q7TFA0 | NS8A_SARS | ORF8a protein | 39 | 27 | 69% |
| **MERS** |  |  |  |  |  |
| **Entry ID** | **Entry Name** | **Protein Name** | **Length**<br>**(AA number)** | **M/H-CrUPs**<br>**(number)** | **M/H-CrUPs**<br>**Density** |
| K9N7C7 | R1AB_MERS1 | Replicase polyprotein 1ab | 7.078 | 5.364 | 76% |
| K9N638 | R1A_MERS1 | Replicase polyprotein 1a | 4.391 | 3.338 | 76% |
| K9N5Q8 | SPIKE_MERS1 | Spike glycoprotein | 1.353 | 1.024 | **76%** |
| K9N4V7 | NCAP_MERS1 | Nucleoprotein | 411 | 301 | 73% |
| K9N643 | ORF4B_MERS | Non-structural protein ORF4b | 246 | 185 | 75% |
| K9N7D2 | ORF5_MERS1 | Non-structural protein ORF5 | 224 | 169 | 75% |
| K9N7A1 | VME1_MERS1 | Membrane protein | 219 | 158 | 72% |
| K9N4V0 | ORF4A_MERS1 | Non-structural protein ORF4a | 109 | 77 | 71% |
| K9N796 | ORF3_MERS1 | Non-structural protein ORF3 | 103 | 74 | 72% |
| K9N5R3 | VEMP_MERS1 | Envelope small membrane protein | 82 | 59 | 72% |

The analysis of the SARS-CoV-2, SARS-CoV and MERS-CoV virus is presented. All the proteins viruses were analyzed and each protein is shown by its EntryID, Entry Name and Protein Name according to the UNIPROT data base. The AA length of each protein and the CrUPs of each protein against the human proteome are shown. Density is defined as the percentage of the total Amino Acids contained in CrUPs of each protein to the total number of the protein's Amino Acids.

In order to illuminate the mechanisms orchestrating the differential pathologies of SARS-CoV-2 compared to other coronavirus family members, we, next, applied the same strategy to other two similar viruses; the Severe Acute Respiratory Syndrome CoronaVirus (SARS-CoV) and the Middle East Respiratory Syndrome-related CoronaVirus (MERS-CoV). Among human viruses, SARS-CoV-2 (C) together with SARS-CoV (S) and MERS-CoV (M) constitute the β coronavirus group and they use the same cellular receptor, the Angiotensin-Converting Enzyme 2 (ACE2), with SARS-CoV-2 sharing approximately 80% and 70% amino acid sequence identity with SARS-CoV and MERS-CoV, respectively (Saputri et al., 2020; Walls et al., 2020). SARS-CoV viral proteome includes 15 reviewed proteins, while MERS-CoV contains 10 reviewed proteins in the UNIPROT database. Our findings confirm the strong similarities among these three coronaviruses at the level of CrUP structure and architecture against human proteome. Interestingly, a more comprehensive analysis of CrUPs per protein has revealed significant differences between them. Intriguingly, the density of M/H-CrUPs per protein ranges between 71-76% (5% range), the density of S/H-CrUPs per protein varies between 61-76% (15% range) and the density of C/H-CrUPs per protein fluctuates between 56-78% (22% range) (Table 2), thus indicating the comparatively more heterogenous CrUPs density in the SARS-CoV-2 coronaviral proteome.

Among all SARS-CoV-2 proteins, the SPIKE_SARS2 (P0DTC2) one (Spike) has received the greatest attention as a key element for virus attachment to the host cell, and as such it has become a principal target for therapeutic vaccine development (Papa et al., 2021; Xia 2021). To mechanistically couple protein’s molecular features with virus pathology at the level of C/H-CrUPs, we comparatively analyzed the Spike proteins of the three coronaviruses and, then, projected the findings onto SPIKE_SARS2 mutation map. Spike glycoprotein presents a length of 1.273 amino acids in SARS-CoV-2, 1.275 amino acids in SARS-CoV and 1.373 amino acids in MERS-CoV (Agrawal et al., 2021). Their densities in CrUPs against the human proteome are measured as 78%, 76% and 76%, respectively, exhibiting the highest CrUP density values among all proteins for each virus herein studied (Table 2). Amino acid sequence alignment of SPIKE_SARS2 (P0DTC2), SPIKE_SARS (P59594) and R9UQ53_MERS (R9UQ53) proved that these three viral Spike proteins share a group of 12 regions, herein defined as Universal Peptides (Fig. S2 and Table S1). The majority of coronaviral Universal Peptides are clustered in the S2 domain of each Spike protein, with a critical one of them (UPs) containing the Furin cleavage site 3 (R^815^↓S).

Most importantly, SARS-CoV-2 Spike protein has presented a significant mutational diversity (Sanches et al., 2021; Tzou et al., 2020). Hitherto, 9 main variants with adaptive mutations and high spread to human populations, named from Alpha to Lambda, respectively, have been thoroughly mapped and characterized. These 8 variants are divided in 39 sub-variants, while other 32 sporadic variants have also been described (Tzou et al., 2020). To investigate the association of mutational profiling with C/H-CrUP landscaping of SARS-CoV-2 Spike protein, the 39 sub-variants together with the wild-type Spike protein (SPIKE_SARS2, P0DTC2) were suitably aligned (Fig S3A). This multiple alignment illustrates all the herein identified Universal Peptides (Table S2) and all the mutations previously announced per isolated variant (Fig S3B). Notably, it seems that almost all the hitherto characterized mutations are identified in regions being located outside the Universal Peptides group. Their majority are clustered in the S1 domain of Spike protein, with two critical mutations being detected in the S1-S2 bridge region, at the amino acid residue 681 that resides in proximity to the first cleavage position by Furin protease, in between the 685^th^ and 686^th^ amino acid residue (Fig S3C) (Davidson et al., 2020; Coutard et al., 2020). Remarkably, all the examined mutations herein prove to create new CrUPs against the human proteome compared to the wild-type Spike protein, thus indicating that the mutant virus strains need novel clinical treatments. This is an important finding, since these new C/H-CrUPs do not exist in the human proteome, but are observed exclusively in the mutant virus proteomes, thereby justifying the great attention Alpha, Delta, Kappa, Lambda and Mu variants have recently received at the worldwide level (Tzou et al., 2020). Table 3, lists all the novel C/H-CrUPs being created by the hitherto reported mutations in coronavirus variants. These variants include 25 mutations, which produce 44 new CrUPs against the human proteome. It may be these novel C/H-CrUPs that give rise to formation of new Intrinsically Disordered Regions (IDRs) and Small Linear Motifs (SLiMs) in the SARS-CoV-2 Spike protein mutant versions (van der Lee et al., 2014; Hraber etal., 2020).

**Table 3.** New C/H-CrUPs of SARS-CoV-2 spike protein in Alpha, Delta, Kappa and Lambda variants.

| Mutations position | Mutation | Variant | New C/H-CrUPs first AA position | New C/H-CrUPs |
| --- | --- | --- | --- | --- |
| 19 | T19R | Delta_P0DTC2 | - | - |
| 70 | V70F | Delta_P0DTC2 | 69 | HFSGTN |
|  |  |  | 70 | FSGTNG |
| 75 - 76 | G75V&T76I | Lambda_P0DTC2 | 71 | SGTNVI |
|  |  |  | 75 | VIKRFD |
| 222 | A222V | Delta_P0DTC2 | 218 | QGFSVL |
| 258 | W258L | Delta_P0DTC2 | - | - |
| 417 | K417N | Delta_P0DTC2 | 413 | GQTGNI |
|  |  |  | 414 | QTGNIA |
| 452 | L452R | Delta_P0DTC2 | 449 | YNYRY |
|  |  | Kappa_P0DTC2 |  |  |
|  |  | Alpha_P0DTC2 |  |  |
|  | L452Q | Lambda_P0DTC2 | 448 | NYNYQ |
|  |  |  | 449 | YNYQY |
| 478 | T478K | Delta_P0DTC2 | 474 | QAGSKP |
|  |  |  | 478 | KPCNG |
| 484 | E484Q | Kappa_P0DTC2 | 481 | NGVQG |
|  |  |  | 483 | VQGFN |
|  |  |  | 484 | QGFNC |
|  | E484K | Alpha_P0DTC2 | 484 | KGFNC |
| 490 | F490S | Lambda_P0DTC2 | 487 | NCYSP |
| 494 | S494P | Alpha_P0DTC2 | - | - |
| 501 | N501Y | Alpha_P0DTC2 | 498 | QPTY |
|  |  |  | 499 | PTYG |
|  |  |  | 500 | TYGV |
|  |  |  | 501 | YGVG |
| 570 | A570D | Alpha_P0DTC2 | 568 | DIDDTT |
| 614 | D614G | Delta_P0DTC2 | 609 | AVLYQG |
|  |  | Kappa_P0DTC2 |  |  |
|  |  | Alpha_P0DTC2 |  |  |
|  |  | Lambda_P0DTC2 |  |  |
|  |  | Delta_P0DTC2 | 610 | VLYQGV |
|  |  | Kappa_P0DTC2 |  |  |
|  |  | Alpha_P0DTC2 |  |  |
|  |  | Lambda_P0DTC2 |  |  |
| 681 | P681R | Delta_P0DTC2 | 680 | SRRRARS |
|  |  | Kappa_P0DTC2 |  |  |
|  | P681H | Alpha_P0DTC2 | 677 | QTNSH |
|  |  |  | 678 | TNSHR |
|  |  |  | 680 | SHRRAR |
| 716 | T716I | Alpha_P0DTC2 | 714 | IPINF |
| 859 | T859N | Lambda_P0DTC2 | 855 | FNGLNV |
|  |  |  | 857 | GLNVLP |
| 950 | D950N | Delta_P0DTC2 | 946 | GKLQN |
|  |  |  | 947 | KLQNVV |
|  |  |  | 948 | LQNVVN |
|  |  |  | 949 | QNVVNQ |
| 982 | S982A | Alpha_P0DTC2 | 978 | NDILAR |
| 1071 | Q1071H | Kappa_P0DTC2 | 1067 | YVPAH |
|  |  |  | 1069 | PAHEKN |
|  |  |  | 1071 | HEKNF |
| 1118 | D1118H | Alpha_P0DTC2 | 1113 | QIITTH |
|  |  |  | 1115 | ITTHN |
|  |  |  | 1116 | TTHNT |
|  |  |  | 1117 | THNTF |
|  |  |  | 1118 | HNTFV |

The new CrUPs of SARS-CoV-2 spike protein (SPIKE\_SARS2, P0DTC2) across the variants Alpha, Delta, Kappa and Lambda were presented. In the first column the position of the mutation in the spike protein sequence is shown. In the second column the mutation is recorded. In the third column is recorder the SARS-CoV-2 main variant in which the mutation appeared. In the fourth column the position of the first amino acid of the new C/H-CrUP created by the mutation is shown. In the last column the new created C/H-CrUPs by the mutation is recorded. The mutant amino acid in the new C/H-CrUPs is in red color. Mutations that not created new C/H-CrUPs is shown by '-'. Some mutations produce multiple new C/H-CrUPs, while 4 new C/H-CrUPs created in more than one variant.

The molecular mechanism of Spike protein’s proteolytic activation has been shown to play a crucial role in the selection of host species, virus binding to the ACE2 receptor, virus-cell fusion and viral infection of human lung cells (Peacock et al., 2021; Whittaker 2021; Shang et al., 2020a). SPIKE_SARS2 (P0DTC2) contains three cleavage sites; the R^685^↓S and R^815^↓S positions that serve as direct targets of the Furin protease, and the T^696^↓M position that can be recognized by the TMPRSS2 protease (Hoffmann et al., 2020a; Hoffmann et al., 2020b; Takeda, 2021). Analysis of the wild-type C/H-CrUPs and the newly formed, mutation-induced, C/H-CrUPs in Spike protein unveiled that the mutation-driven, novel, peptides are created exclusively, around the R^685^↓S cleavage site, by the two pathogenic mutations P681H and P681R (Table S2).

Notably, among these four new peptides, the only one new peptide that embraces Furin’s cleavage site is the “SRRRAR↓S” C/H-CrUP, which is solely generated by the P681R mutation carried by the Delta and Kappa coronavirus variants, while at the same time the peptide “PRRARSV” conserve its uniqueness even after the replacement of Proline with Arginine and its transformation to “RRRARSV” (Fig. 2).

**Figure 2.**
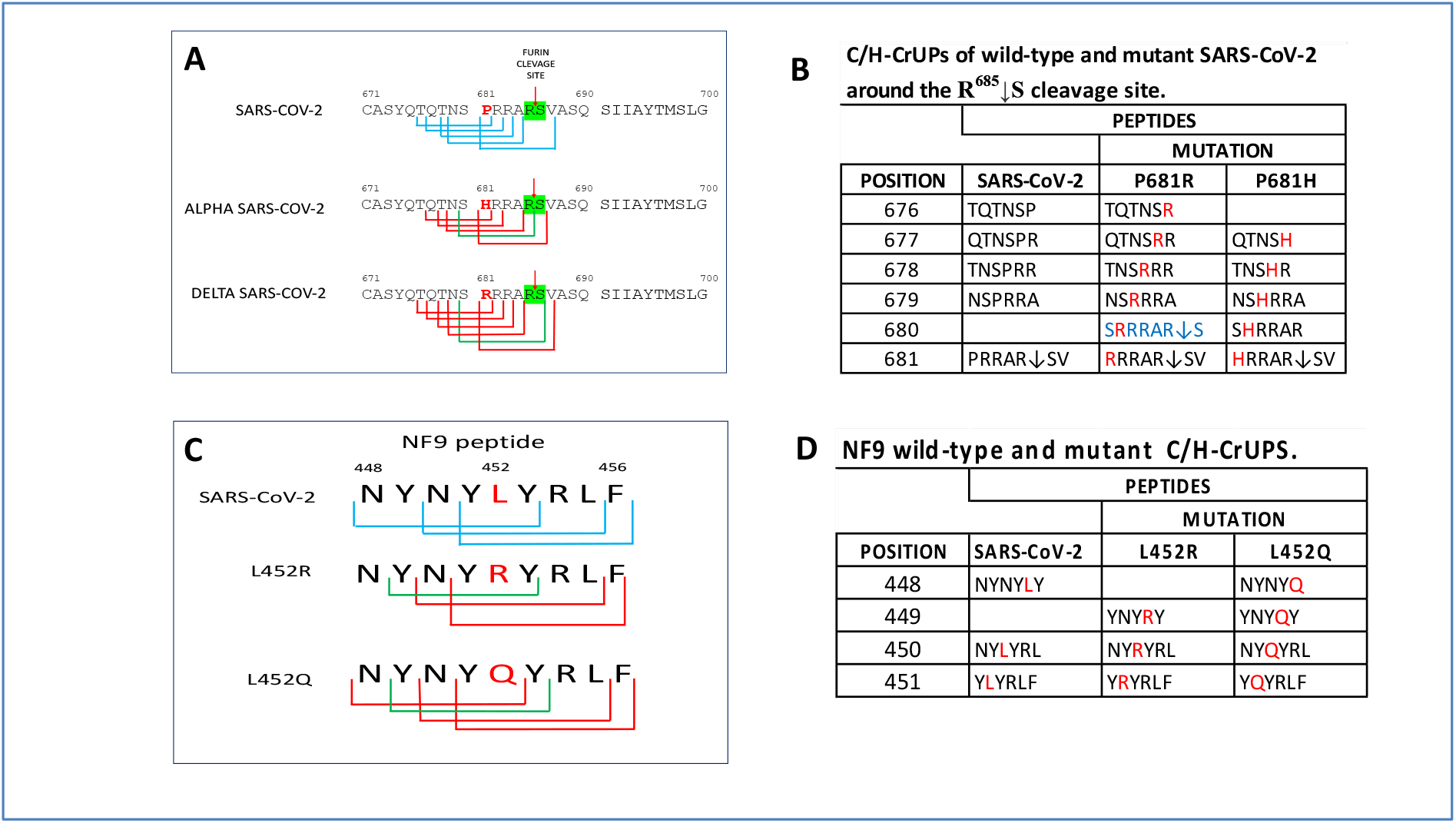
C/H- CrUPs around the R^685^↓S cleavage site and of NF9 peptide of SPIKE_SARS2. **A)** The sequence of the wild-type SPIKE_SARS2 protein between the position 671 up to 700 in wild-type, Alpha and Delta variants of SARS-CoV-2, spike protei is shown. In each variant the C/H-CrUPs were marked. Blue lines indicate the C/H-UPs of wild-type protein around the R^685^↓S cleavage site. Red lines indicate the C/H-CrUPs resulted by the P681H and P681R mutations. Green Lines the new created mutant C/H-CrUPs by the P681H and P681R mutations in Alpha and Delta variant respectively. **B)** Table of C/H-CrUPs around the R^685^↓S of the wild-type and mutant spike protein. **C)** The sequence of the wild-type NF9 protein between the position 448 up to 456 in wild-type SPIKE_SARS2 protein and after the L452R and L452Q mutations. Blue lines indicate the C/H-UPs of NF9 peptide. Red lines indicate the C/H-CrUPs resulted by the L452R and L452Q mutations. Green Lines the new created mutant C/H-CrUPs by the L452R and L452Q mutations. **D)** Table of NF9 peptide C/H-CrUPs in wild-type and after the L452R and L452Q mutation.

The Furin cleavage site R^685^↓S has been characterized as a 20 amino acid motif that corresponds to the amino acid sequence A672-S691 of the SPIKE_SARS2 (P0DTC2) protein (Fig 2) (Wu and Zhao, 2020). The 8 amino acid sequence peptide “SPRRAR↓SV” (S680-V687) serves as the core region of the motif, while two flanking solvent-accessible regions of 8 amino acids (A672-N679) and 4 amino acids (A688-S691) long, respectively, are recognized (Takeda, 2021; Wu and Zhao, 2020).

Pro-protein Convertase (PC) Furin and/or Furin-like PCs act as sequence-specific proteases, and can cleave the Spike protein in a position recognizing the unique, and positively charged by the Arginine, motif “R-x-x-R↓S” (Wu and Zhao, 2020). Since Furin and/or Furin-like PCs are secreted from host cells and bacteria in the airway epithelium, while other PCs, such as the PC5/6A and PACE4, exhibit widespread tissue distribution, it is likely that their activities may be critically implicated in the SARS-CoV-2-induced damage and pathology of multiple infected organs (Örd et al., 2020). It seems that Furin’s cleavage site essentially contributes to the infection process and disease progression, and offers a powerful target for immunogenetic, antigenic and therapeutic interventions, as corroborated by the recently developed new antibody against Furin’s cleavage site (Braun et al., 2019; Zahradník et al., 2021; Wu et al, 2020).

Most importantly, the SARS-CoV-2 Delta variant that carries the critical mutation P681R seems to be more infectious and pathogenic than the wild-type virus form, while the importance of that mutation has very recently begun to be recognized (Wu et al., 2020). Replacement of Proline with Arginine at position 681 causes the loss of amino acid sequence uniqueness that characterizes the wild-type “PRRARSV” C/H-CrUP and likely increases the possibility of Furin’s cleavage site (core region) to be significantly stabilizing its conformation, thus facilitating a more efficient Spike protein cleavage process by the Furin protease (Whittaker, 2021; Callaway, 2021). To the same direction, novel SLiMs, such as “SRRR”, “RRR”, “RRRAR” and “RRRARS”, can be produced by the mutant C/H-CrUPs, which may act as specific targets of other than Furin PCs, thereby enabling the stronger (and quicker) binding of the mutant virus to its host ACE2 receptor that likely leads to a comparatively more generalized infection and massive mutant virus production (Table S3) (Shorthouse et al.,2021; Davey et al., 2015). That fact seems to be evident by the dramatic increase of the total number of motifs created by the P681R mutation identified within the Human proteome (Table S3). Of note, the mutant C/H-CrUP-derived new SLiMs, in the SARS-CoV-2 Delta variant, could render Spike protein antigenically weak or defective, fostering it to lose its capacity to serve as antibody target promotes the virus immune escape (Davey et al., 2015; Almehdi et al., 2021).

An important issue for viral infectivity and pathogenesis is the receptor recognition and binding of the virus to the host cell surface. SARS-CoV-2 belongs to the β coronavirus group and, like SARS-CoV, uses the same cellular receptor, the Angiotensin-Converting Enzyme 2 (ACE2) (Walls et al., 2020; Wang et al., 2020). The SARS-CoV-2 Spike protein attaches to ACE2 receptor by a Receptor-Binding Domain (RBD) defined in the Spike protein from positions F318 up to F541 (Shang et al., 2020b). Nowadays, this region has received great attention, as it seems to be the target of antibodies against the virus and other therapeutic interventions (Chen et al., 2021; Zahradník et al., 2021; Hastie et al., 2021). Additional studies have shown that from the amino acid residue W436 up to the Q506 one the RBD contains the Receptor-Binding Motif (RBM), which carries 12 contact positions with ACE2 (Hatmal et al., 2020). Mutation analysis revealed that in 10 positions of the RBD region 13 mutations were described (Fig. S3 and Table S4). In RBM, 10 mutations in 6 sequence positions were described in different virus variants (Table S4), while from the 10 contact positions only the P501Y in Alpha, Beta, Gamma and Mu variants was found to be mutated (Table S5).

The most important region in RBM is the peptide NYNYLYRLF (from 448 to 456 position). This tyrosine-enriched peptide contains two contact site (Y449 and Y453) and is known as the NF9 peptide (Motozono et al., 2021). It seems to affect antigen recognition, by being an immunodominant HLA*24:02-restricted epitope identified by CD8^+^ T cells. Furthermore, NF9 stimulation also increases cytokine production produced from CD8^+^ T cells, such as IFN-γ, TNF-α and IL-2 (Kared et al., 2021). Analysis of C/H-CrUPs of the NF9 peptide showed that it contains 3 unique peptides (Fig. 2D and E, and Table S6). Mutation analysis indicated that in the NF9 peptide the mutation L452R occurs in the variants Alpha, Delta, Iota and Kappa, while the mutation L452Q appears in the variant Lambda. Further analysis unveiled that these mutations are observed in the amino acid that resides at position 5, exactly in the middle of the peptide, creating 3 and 4 new C/H CrUPs, respectively (Table S6). These mutations have a dramatic effect in the uniqueness of the NF9 peptide(s). Namely, the 6 amino acid length C/H-CrUPs “NYNYLY” losses its uniqueness against the human proteome, while only by the mutation L452Q a new core unique peptide with 5 amino acid length is surprisingly created (Fig. 2D and E, and Table S6). The loss of uniqueness of this peptide, which notably is located at the beginning of NF9 peptide, seems to be crucial, as it leads to the loss of the antigenic capacity of the NF9 peptide, thus evading the HLA-A24-restricted immunity and inducing the immune escape of the virus. Interestingly, related studies have shown that the L452R mutation (and subsequently the newly created C/H-CrUPs herein characterized) increases the infectiveness of SARS-CoV-2, by strengthening the electrostatic interactions of this region on Spike protein with the ACE2 virus receptor (Motozono et al., 2021). Hitherto, epidemiological data indicated that the dominant variant of SARS-CoV-2 is the Delta variant (Mlcochova et al., 2021). Under the light of the aforementioned findings, variant’s enhanced pathogenicity seems to be the outcome of the simultaneous existence (accumulation) of two critical mutations; the L452R and P681R ones, in Delta variant. The mutation L452R, through the loss of NF9 peptide uniqueness, causes virus immune escape and stronger binding of the virus to its cognate receptor, while at the same time mutation P681R facilitates the Spike protein cleavage process by different proteases, inducing a generalized infection and a massive virus release. Therefore, the Delta variant gains a significant advantage of escape from the immune system *per se*, as well as from the vaccination-induced immunity, together with an increased infectiveness, as a result of virus entrance into the host cell, and an increase of virus formation and its massive release.

Interestingly, although mutations outside the Spike protein locus in SARS-CoV-2 coronavirus genome have not been yet completely mapped, in a systematic manner, our study also reveals novel and useful information of all the remaining (Spike protein-independent) C/H-CrUPs that seem to hold strong promise and open a new therapeutic window for the Covid-19 pandemic. Finally, the approach of virus-host unique peptide signature identification could prove a useful tool for the elucidation of virus infectiveness, prevention of virus immune escape, domination of pathogenic variants, and identification of new antigenic and pharmacological targets.

## Supporting information

Pierros et al. Suppl. Material

## Author contributions

Conceptualization: VP, EK, GThT, Methodology: VP, EK, Investigation: VP, EK, GThT, Visualization: EK, DJS, GThT, Supervision: GThT, Writing – original draft: DJS, GThT, Writing – review & editing: DJS, GThT

## Data and materials availability

All data are available in the main text or the supplementary materials

## Methods

A new bioinformatic tool was developed which can extract from a given proteome the Core Unique Peptides (thus creating it’s Uniquome). The user can specify the min and max peptide lengths that the tool will analyze. The tool will split each protein to all possible peptides of length min to length max thus generating a very large set of peptides (for a protein of length **L** with a window of size **W** a set of **C = L -W + 1** will be generated). In the next step all these peptides starting from smallest to largest will be searched against the rest of the proteome to decide whether the peptide exists on another protein or not. Since the search is for the smallest possible peptide (Core Unique Peptide) the tool will first make sure that the peptide under examination does not already contain a smaller Core Unique Peptide. This is ensured by examining if any of the already identified Core Unique Peptides of the protein is contained within the peptide under examination. All peptides that conform to these 2 rules are **Core Unique Peptides**.

The following diagram describes the algorithm we use to identify these Core Unique Peptides.

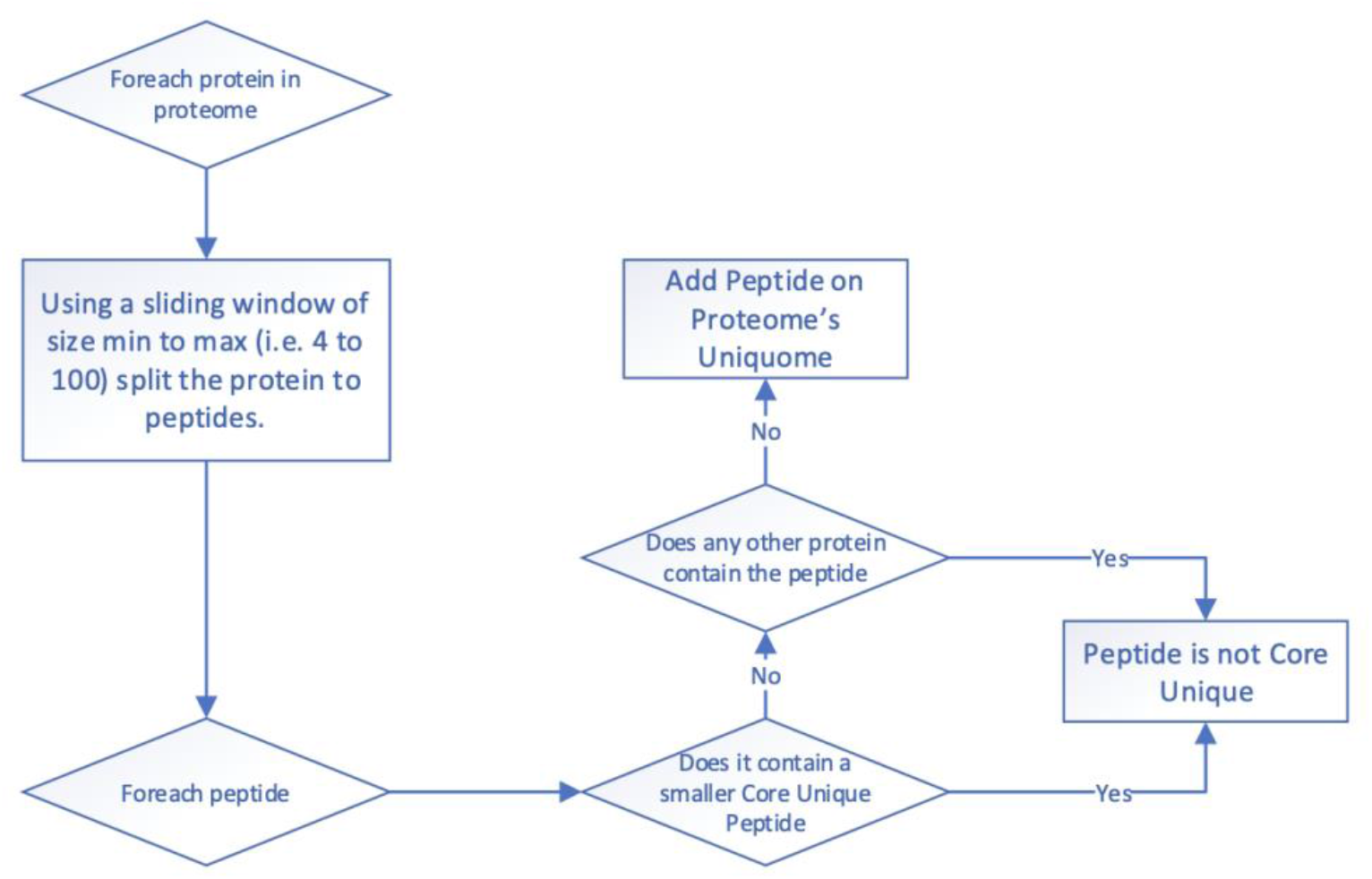

In the following figure a sliding window of 9 aminoacids is applied on O00400 ACATN_HUMAN protein generating candidate peptides VYVKNFGRR and YVKNFGRRK. Those peptides will be searched against the rest of the proteome to determine their uniqueness once we have determined that they don’t already contain a smaller Core Unique Peptide. The latter is determined by examining whether an already defined Core Unique Peptide is contained within the peptide.

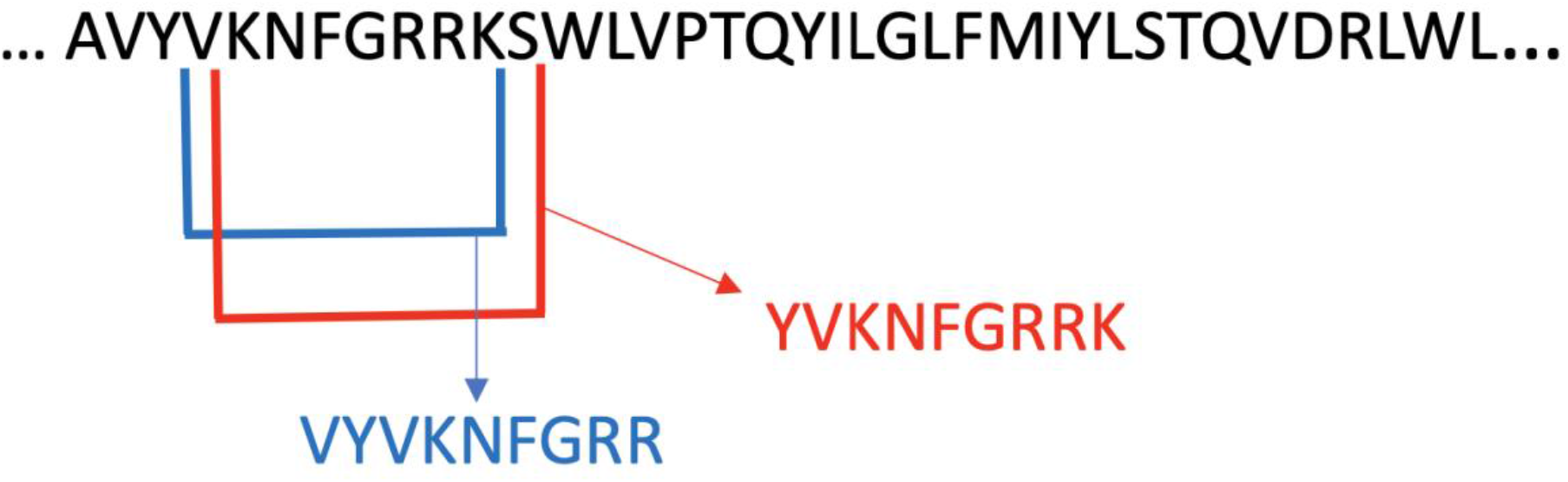

To address the hypothesis of the current study, the aforementioned tool was expanded by developing a new feature where the user can give a reference and a target proteome. This new feature allows the tool to search all the peptides of the target proteome against the reference proteome thus creating a set of Core unique peptides of **Target** vs **Reference** proteomes. To accomplish that the tool will (like on the initial implementation) split all proteins in the target proteome to all possible peptides of length min to length max. Now instead of searching for the uniqueness of each peptide within the same proteome, it performs that search against the **reference** proteome. Like before the peptide under examination must not contain any smaller peptides already identified as **Core Unique Peptides**.

The following diagram describes the algorithm we use to identify these Core Unique Peptides.

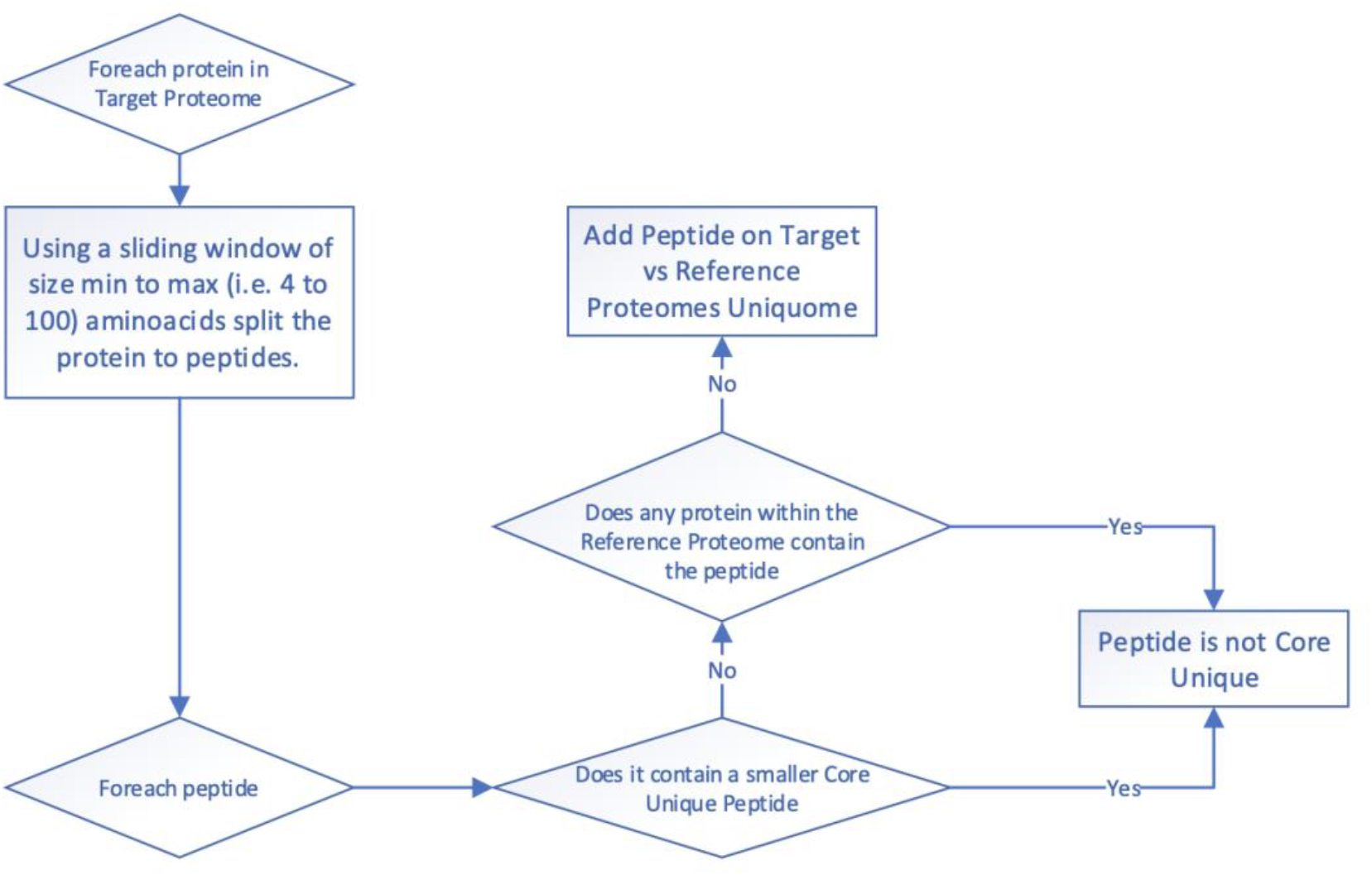

## Motifs and SLiMs search

For Motifs and SLiMs identification and search, the tool offers the user the ability to perform a motif search to identify possible SLiMs. User gives an N length peptide as well as the number of aminoacids that can vary in the given peptide. The tool then creates all possible combinations of peptides that can be produced by considering in each combination exactly N aminoacid(s) as unknown. Once those combinations are produced an exhaustive search using regular expressions is performed against the **reference proteome** to locate all possible proteins containing such peptides. To better highlight the process, if the user provides the peptide **TQYILG** and **N=2** the following combinations will be produced:

- **??YILG**
- **?Q?ILG**
- **?QY?LG**
- **?QYI?G**
- **?QYIL?**
- **T??ILG**
- **T?Y?LG**
- **T?YI?G**
- **T?YIL?**
- **TQ??LG**
- **TQ?I?G**
- **TQ?IL?**
- **TQY??G**
- **TQY?L?**
- **TQYI??**

User will receive a list of all the proteins which contain peptides that matches the criteria including the motif against which the peptide was matched and all the positions within the protein sequence where that peptide can be found. All proteomes were taken from Uniprot

## Data bases

All proteomes and proteins were obtained from: Uniprot [https://www.uniprot.org].

SARS-Cov_2 wild type, variants sequences and mutations were obtained from Stanford COVID Database [https://covdb.stanford.edu/page/mutation-viewer/]

Motifs were taken from the Eukaryotic Linear Motif resource for Functional Sites in Proteins [http://elm.eu.org/index.html] and KEGG/GenomeNet/MOTIF2 [https://www.genome.jp/tools/motif/MOTIF2.html]

SLiMs containing proteins were taken from Davey lab SLiMs servers (The Institute of Cancer Research, UK (ICR) [http://slim.icr.ac.uk/slimsearch/] and [http://slim.icr.ac.uk/index.php?page=tools]

