## Supplementary material for "Unique Peptide Signatures Of SARS-CoV-2 Against Human Proteome Reveal Variants’ Immune Escape And Infectiveness": Pierros et al. Suppl. Material

### SUPPLEMENTARY MATERIALS

**Suppl. Figures 1-3**
**Suppl. Tables 1-6**

A

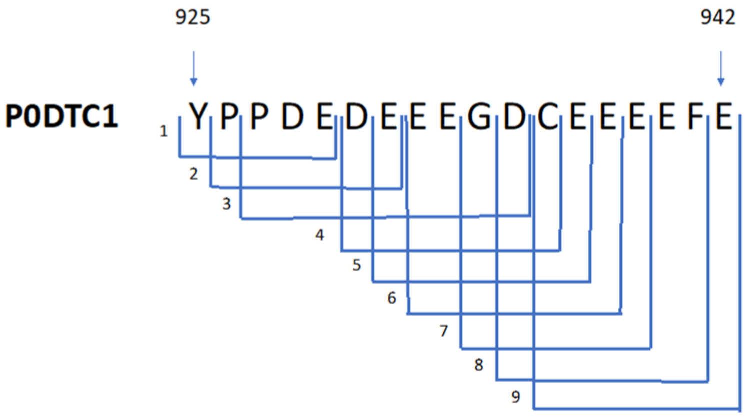


B

**C/H-CrUPs identified from the position 925 to 942 of R1A_SARS2 (P0DTC1) protein**

| Peptide Number | AA sequence | Potition | AA number |
|---|---|---|---|
| 1 | YPPDE | 925-929 | 5 |
| 2 | PPDEDE | 926-931 | 6 |
| 3 | PDEDEEEGD | 927-935 | 9 |
| 4 | DEEEGDC | 930-936 | 7 |
| 5 | EEEGDCE | 931-937 | 7 |
| 6 | EEGDCEE | 932-938 | 7 |
| 7 | GDCEEE | 934-939 | 6 |
| 8 | DCEEEEF | 935-941 | 7 |
| 9 | CEEEEFE | 936-942 | 7 |

**Suppl. Fig. 1. Identification of C/H-CrUPs around the position of AA925-AA942 of the SARS-CoV-2 protein R1A_SARS2 (P0DTC1).** Between these positions one of the two C/H-CrUPs with a 9AA length is included from the position 927-935. **A)** Schematic representation of the 9 C/H-CrUPs peptides included in that peptide, **B)** Table of C/H-CrUPs.

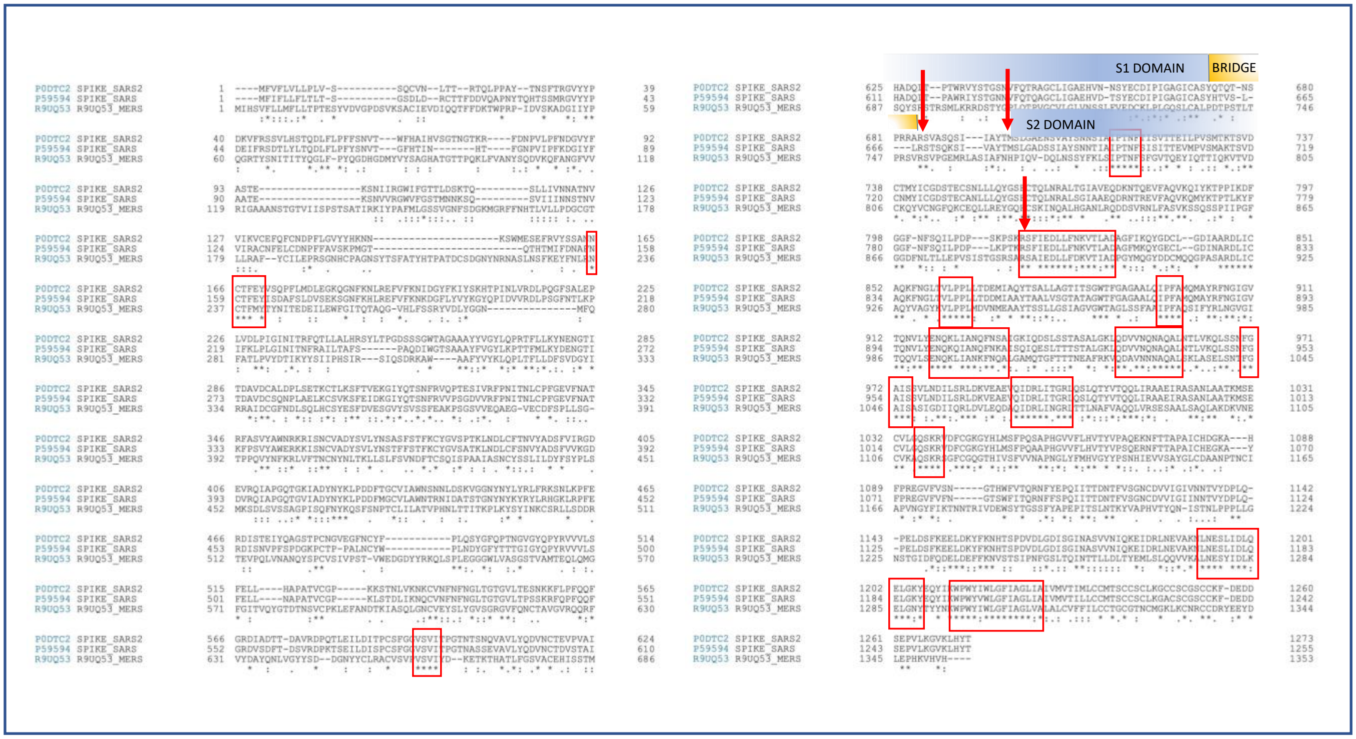


**Suppl. Fig. 2. Alignment of the SARS-CoV-2, SARS-CoV and MERS-CoV spike proteins.** The sequence of the spike proteins P0DTC2, P59594 and R9UQ53 of the above viruses respectively were obtained for Uniprot Data base and alignment according to an available application in that database. Green blocks with red outline mark the identical peptidic sequences between the alignment sequences. These identical peptidic sequences were considered as Universal Peptides. Red arrows mark the cleavage sites of the SARS-CoV-2 SPIKE_protein

A

START — SIGNAL PEPTIDE … END

| Variant | 1 | 2 | 3 | 4 | 5 | 6 | 7 | 8 | 9 | 10 | 11 | 12 | 13 | 1260 | 1261 | 1262 | 1263 | 1264 | 1265 | 1266 | 1267 | 1268 | 1269 | 1270 | 1271 | 1272 | 1273 |
|---|---|---|---|---|---|---|---|---|---|---|---|---|---|---|---|---|---|---|---|---|---|---|---|---|---|---|---|
| SARS-CoV-2 wildtype | M | F | V | F | L | V | L | L | P | L | V | S | S | D | S | E | P | V | L | K | G | V | K | L | H | Y | T |
| Alpha (B.1.1.7 & Q1-Q4) | M | F | V | F | L | V | L | L | P | L | V | S | S | D | S | E | P | V | L | K | G | V | K | L | H | Y | T |
| Alpha (B.1.1.7+E484K) | M | F | V | F | L | V | L | L | P | L | V | S | S | D | S | E | P | V | L | K | G | V | K | L | H | Y | T |
| Alpha (B.1.1.7+L452R) | M | F | V | F | L | V | L | L | P | L | V | S | S | D | S | E | P | V | L | K | G | V | K | L | H | Y | T |
| Alpha (B.1.1.7+S494P) | M | F | V | F | L | V | L | L | P | L | V | S | S | D | S | E | P | V | L | K | G | V | K | L | H | Y | T |
| Beta (B.1.351) | M | F | V | F | L | V | L | L | P | L | V | S | S | D | S | E | P | V | L | K | G | V | K | L | H | Y | T |
| Beta (B.1.351+E516Q) | M | F | V | F | L | V | L | L | P | L | V | S | S | D | S | E | P | V | L | K | G | V | K | L | H | Y | T |
| Beta (B.1.351+L18F & B.1.351.2-3) | M | F | V | F | L | V | L | L | P | L | V | S | S | D | S | E | P | V | L | K | G | V | K | L | H | Y | T |
| Beta (B.1.351+P384L) | M | F | V | F | L | V | L | L | P | L | V | S | S | D | S | E | P | V | L | K | G | V | K | L | H | Y | T |
| Gamma (P.1 & P.1.1 - P.1.2) | M | F | V | F | L | V | L | L | P | L | V | S | S | D | S | E | P | V | L | K | G | V | K | L | H | Y | T |
| Gamma (P1+P681H) | M | F | V | F | L | V | L | L | P | L | V | S | S | D | S | E | P | V | L | K | G | V | K | L | H | Y | T |
| Delta (B.1.617.2) | M | F | V | F | L | V | L | L | P | L | V | S | S | D | S | E | P | V | L | K | G | V | K | L | H | Y | T |
| Delta (B.1.617.2+E484Q) | M | F | V | F | L | V | L | L | P | L | V | S | S | D | S | E | P | V | L | K | G | V | K | L | H | Y | T |
| Delta (B.1.617.2+K417N) | M | F | V | F | L | V | L | L | P | L | V | S | S | D | S | E | P | V | L | K | G | V | K | L | H | Y | T |
| Delta (B.1.617.2+Q613H) | M | F | V | F | L | V | L | L | P | L | V | S | S | D | S | E | P | V | L | K | G | V | K | L | H | Y | T |
| Delta (AY.1) | M | F | V | F | L | V | L | L | P | L | V | S | S | D | S | E | P | V | L | K | G | V | K | L | H | Y | T |
| Delta (AY.2) | M | F | V | F | L | V | L | L | P | L | V | S | S | D | S | E | P | V | L | K | G | V | K | L | H | Y | T |
| Delta (AY.3 - AY.8 & AY.12) | M | F | V | F | L | V | L | L | P | L | V | S | S | D | S | E | P | V | L | K | G | V | K | L | H | Y | T |
| Delta (AY.9) | M | F | V | F | L | V | L | L | P | L | V | S | S | D | S | E | P | V | L | K | G | V | K | L | H | Y | T |
| Delta (AY.10) | M | F | V | F | L | V | L | L | P | L | V | S | S | D | S | E | P | V | L | K | G | V | K | L | H | Y | T |
| Delta (AY.11) | M | F | V | F | L | V | L | L | P | L | V | S | S | D | S | E | P | V | L | K | G | V | K | L | H | Y | T |
| Eta (B.1.525) | M | F | V | F | L | V | L | L | P | L | V | S | S | D | S | E | P | V | L | K | G | V | K | L | H | Y | T |
| Iota (B.1.526) | M | F | V | F | F | V | L | L | P | L | V | S | S | D | S | E | P | V | L | K | G | V | K | L | H | Y | T |
| Kappa (B.1.617.1) | M | F | V | F | L | V | L | L | P | L | V | S | S | D | S | E | P | V | L | K | G | V | K | L | H | Y | T |
| Lambda (C.37) | M | F | V | F | L | V | L | L | P | L | V | S | S | D | S | E | P | V | L | K | G | V | K | L | H | Y | T |
| Mu (B.1.621) | M | F | V | F | L | V | L | L | P | L | V | S | S | D | S | E | P | V | L | K | G | V | K | L | H | Y | T |

POINT MUTATION

SPIKE UNIVERSAL PEPTIDES

B

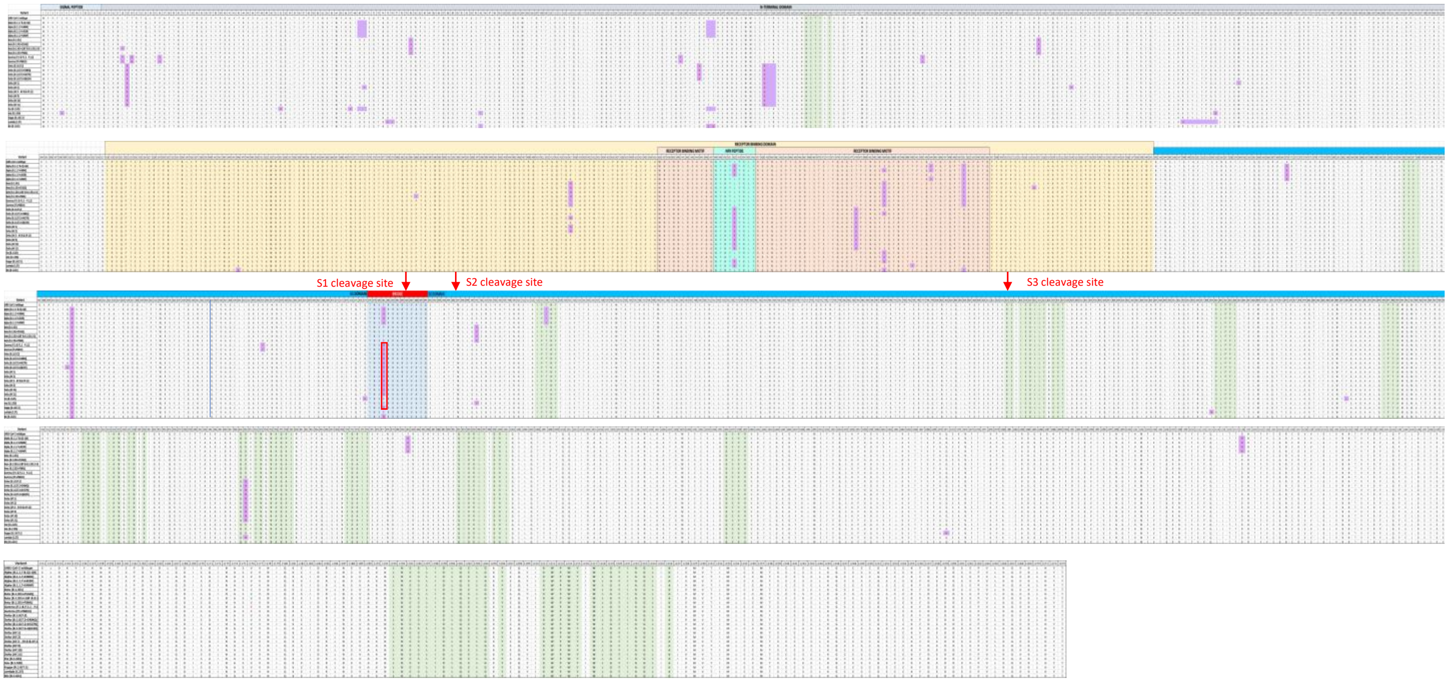


POINT MUTATION

UNIVERSAL PEPTIDES

FURIN CLEVAGE SITE

C

| | S1 DOMAIN | | | | | | | | BRIDGE | | | | | | | | | | | | | S2 DOMAIN | | | | |
|---|---|---|---|---|---|---|---|---|---|---|---|---|---|---|---|---|---|---|---|---|---|---|---|---|---|---|
| **Variant** | 670 | 671 | 672 | 673 | 674 | 675 | 676 | 677 | 678 | 679 | 680 | 681 | 682 | 683 | 684 | 685 | 686 | 687 | 688 | 689 | 690 | 691 | 692 | 693 | 694 | 695 |
| SARS-CoV-2 wildtype | I | C | A | S | Y | Q | T | Q | T | N | S | P | R | R | A | R | S | V | A | S | Q | S | I | I | A | Y |
| Alpha (B.1.1.7 & Q1-Q4) | I | C | A | S | Y | Q | T | Q | T | N | S | H | R | R | A | R | S | V | A | S | Q | S | I | I | A | Y |
| Alpha (B.1.1.7+E484K) | I | C | A | S | Y | Q | T | Q | T | N | S | H | R | R | A | R | S | V | A | S | Q | S | I | I | A | Y |
| Alpha (B.1.1.7+L452R) | I | C | A | S | Y | Q | T | Q | T | N | S | H | R | R | A | R | S | V | A | S | Q | S | I | I | A | Y |
| Alpha (B.1.1.7+S494P) | I | C | A | S | Y | Q | T | Q | T | N | S | H | R | R | A | R | S | V | A | S | Q | S | I | I | A | Y |
| Beta (B.1.351) | I | C | A | S | Y | Q | T | Q | T | N | S | P | R | R | A | R | S | V | A | S | Q | S | I | I | A | Y |
| Beta (B.1.351+E516Q) | I | C | A | S | Y | Q | T | Q | T | N | S | P | R | R | A | R | S | V | A | S | Q | S | I | I | A | Y |
| Beta (B.1.351+L18F & B.1.351.2-3) | I | C | A | S | Y | Q | T | Q | T | N | S | P | R | R | A | R | S | V | A | S | Q | S | I | I | A | Y |
| Beta (B.1.351+P384L) | I | C | A | S | Y | Q | T | Q | T | N | S | P | R | R | A | R | S | V | A | S | Q | S | I | I | A | Y |
| Gamma (P.1 & P.1.1 - P.1.2) | I | C | A | S | Y | Q | T | Q | T | N | S | H | R | R | A | R | S | V | A | S | Q | S | I | I | A | Y |
| Gamma (P1+P681H) | I | C | A | S | Y | Q | T | Q | T | N | S | H | R | R | A | R | S | V | A | S | Q | S | I | I | A | Y |
| Delta (B.1.617.2) | I | C | A | S | Y | Q | T | Q | T | N | S | R | R | R | A | R | S | V | A | S | Q | S | I | I | A | Y |
| Delta (B.1.617.2+E484Q) | I | C | A | S | Y | Q | T | Q | T | N | S | R | R | R | A | R | S | V | A | S | Q | S | I | I | A | Y |
| Delta (B.1.617.2+K417N) | I | C | A | S | Y | Q | T | Q | T | N | S | R | R | R | A | R | S | V | A | S | Q | S | I | I | A | Y |
| Delta (B.1.617.2+Q613H) | I | C | A | S | Y | Q | T | Q | T | N | S | R | R | R | A | R | S | V | A | S | Q | S | I | I | A | Y |
| Delta (AY.1) | I | C | A | S | Y | Q | T | Q | T | N | S | R | R | R | A | R | S | V | A | S | Q | S | I | I | A | Y |
| Delta (AY.2) | I | C | A | S | Y | Q | T | Q | T | N | S | R | R | R | A | R | S | V | A | S | Q | S | I | I | A | Y |
| Delta (AY.3 - AY.8 & AY.12) | I | C | A | S | Y | Q | T | Q | T | N | S | R | R | R | A | R | S | V | A | S | Q | S | I | I | A | Y |
| Delta (AY.9) | I | C | A | S | Y | Q | T | Q | T | N | S | R | R | R | A | R | S | V | A | S | Q | S | I | I | A | Y |
| Delta (AY.10) | I | C | A | S | Y | Q | T | Q | T | N | S | R | R | R | A | R | S | V | A | S | Q | S | I | I | A | Y |
| Delta (AY.11) | I | C | A | S | Y | Q | T | Q | T | N | S | R | R | R | A | R | S | V | A | S | Q | S | I | I | A | Y |
| Eta (B.1.525) | I | C | A | S | Y | Q | T | H | T | N | S | P | R | R | A | R | S | V | A | S | Q | S | I | I | A | Y |
| Iota (B.1.526) | I | C | A | S | Y | Q | T | Q | T | N | S | P | R | R | A | R | S | V | A | S | Q | S | I | I | A | Y |
| Kappa (B.1.617.1) | I | C | A | S | Y | Q | T | Q | T | N | S | R | R | R | A | R | S | V | A | S | Q | S | I | I | A | Y |
| Lambda (C.37) | I | C | A | S | Y | Q | T | Q | T | N | S | P | R | R | A | R | S | V | A | S | Q | S | I | I | A | Y |
| Mu (B.1.621) | I | C | A | S | Y | Q | T | Q | T | N | S | H | R | R | A | R | S | V | A | S | Q | S | I | I | A | Y |

**Suppl. Fig. 3. Alignment of the SARS-CoV-2 spike protein (SPIKE_SARS2, P0DTC2) of the 25 sup-variants of the main 9 virus variants, together with the native Spike Protein (SPIKE_SARS2, P0DTC2). A)** The protein's N-terminal and Carboxy-terminal are presented for the native spike protein and the 39 virus sup-variants. **B)** The complete spike sequence alignment. Purple blocks marked the point mutations sites in the variants, green color indicate the Universal Peptides of the spike proteins from Fig. S2. Yellow color mark the Receptor-Binding Domain of spike protein to ACE2, pink color mark the Receptor-Binding Motif, cyan mark the NF9 peptide and light blue mark the Bridge between S1 and S2 domain. Red arrows indicate the cleavage sites. With different colors in the upper side of the alignment, the different domains of the spike protein are marked. **C)** The spike protein alignment around the bridge domain (light blue color) between the S1 domain and the S2 domain is presented. Red arrow indicates the furin cleavage site $R^{685}$↓S. Purple blocks mark the point mutations around that position while red outline indicates the Delta and Kappa variants carrying the mutation P681R.

**Suppl. Table 1. Spike Universal Peptides and their actually included CrUP against the Human proteome..**

| SITE | | | | SEQUENCE | CrUP against human proteome | DOMAIN | |
|---|---|---|---|---|---|---|---|
| 165-168 | 170 | | | N C T F * Y | CTFEY | S1 Domain | N-Terminal domain |
| 595-598 | | | | V S V I | VSVITP | S1 Domain | |
| 714-718 | | | | I P T N F | IPTNFT | S1 Domain | |
| 815-816 | 818-823 | 825-827 | 829-830 | R S * I E D L L F * K V T * A D | RSFIED | S2 Domain | S3 Cleavage site (Furin) |
| | | | | | SFIEDL | | |
| | | | | | FIEDLL | | |
| | | | | | **IEDLLF** | | |
| | | | | | EDLLFN | | |
| | | | | | DLLFNK | | |
| | | | | | LLFNKV | | |
| | | | | | LFNKVT | | |
| 860-864 | | | | V L P P L | VLPPLLT | S2 Domain | |
| 896-899 | | | | I P F A | IPFAMQ | S2 Domain | Internal fusion peptide |
| 918-921 | 923-925 | 927-928 | 930 | E N Q K * I A N * F N * A | ENQKLI | S2 Domain | |
| | | | | | QKLIAN | | |
| | | | | | KLIANQ | | |
| | | | | | LIANQF | | |
| | | | | | IANQFN | | |
| | | | | | ANQFNS | | |
| | | | | | NQFNSA | | |
| 949-950 | 952-953 | 955-959 | | Q D * V N * N A Q A L | DVVNQN | S2 Domain | Heptad Repeats 1 |
| | | | | | VVNQNA | | |
| | | | | | NQNAQA | | |
| | | | | | NAQALN | | |
| 970-974 | | | | F G A I S | FGAISSV | S2 Domain | Heptad Repeats 1 |
| 992-997 | 999-1001 | | | Q I D R L I * G R L | **QIDRLI** | S2 Domain | |
| | | | | | IDRLIT | | |
| | | | | | DRLITG | | |
| | | | | | RLITGR | | |
| 1036-1039 | | | | Q S K R | QSKRVD | S2 Domain | |
| 1193-1204 | 1206 | | | L N E S L I D L Q E L G * Y | **NESLID** | S2 Domain | Heptad Repeats 2 |
| | | | | | **SLIDLQ** | | |
| | | | | | **LIDLQE** | | |
| | | | | | **IDLQELG** | | |
| | | | | | DLQELGK | | |
| | | | | | QELGKY | | |
| 1211-1216 | 1217-1224 | | 1226 | K W P W Y * W L G F I A G L * A | **WPWY** | S2 Domain | Trans membrane domair |
| | | | | | PWYIWL | | |
| | | | | | YIWLG | | |
| | | | | | IWLGF | | |
| | | | | | **WLGFIA** | | |
| | | | | | **LGFIAGL** | | |
| | | | | | FIAGLI | | |
| | | | | | IAGLIA | | |

The Universal peptides of SARS-CoV-2, SARS-CoV and MERS-CoV spike proteins according to Fig. S2 presented alignment. The position in the protein sequence and the peptide sequence are shown. Green blocks indicated the common Amino Acids in all proteins, * indicated positions with different Amino Acids between proteins. Next the CrUP included in the Universal Peptides are recorded followed by the domain of the spike protein in which they belong. Yellow blocks indicated complete sequence CrUP appeared in the Universal peptide in all spike proteins alignment

**Suppl. Table 2. New C/H-CrUPs around the SARS-CoV-2 spike protein cleavage sites.**

| Cleavage site | Mutation | Variant | New C/H-CrUPs first AA position | New C/H-CrUP |
|---|---|---|---|---|
| R$^{685}$↓S | P681R | Delta & Kappa | 680 | SRRRAR↓S |
| | P681H | Alpha & Gamma | 677 | QTNSH |
| | | | 678 | TNSHR |
| | | | 680 | SHRRAR |
| T$^{696}$↓M | A701V | Beta | None | |
| R$^{815}$↓S | None | | None | |

The new C/H-CrUPs created by the mutations around the SARS-CoV-2 spike protein (SPIKE_SARS2, P0DTC2) were identified. Fist column: The cleavage site of SARS-CoV-2 spike protein. ↓ indicates the cleavage site. Second column: The mutation identified around the cleavage site. Third column: The virus variants in which the mutation appeared. Fourth column: The position in the SARS-CoV-2 spike protein sequence in which the first Amino Acid of the C/H-CrUP appeared. Fifth column: The sequence of the new C/H-CrUP. ↓ indicates the cleavage site within that peptide.

**Suppl. Table 3. Small Linear Motifs (SLiMs) of wildtype C/H-CrUPs and C/H-CrUPs created by the mutation P681R detected in Human proteome.**

| Motif | Number of proteins in UNIPROT contain the motif | MOTIF FOUND | PROTEIN Entry ID | PROTEIN Entry Name | PROTEIN full Name |
|---|---|---|---|---|---|
| RRRARSV | 0 | - | - | | |
| XRRARSV | 1 | ARRARSV | P37088 | SCNNA_HUMAN | Amiloride-sensitive sodium channel subunit alpha |
| RXRARSV | 1 | RPRARSV | Q86X29 | LSR_HUMAN | Lipolysis-stimulated lipoprotein receptor |
| RRXARSV | 2 | RRDARSV | Q8WWN8 | ARAP3_HUMAN | Arf-GAP with Rho-GAP domain, ANK repeat and PH domain-containing protein 3 |
| | | RRPARSV | Q9H427 | KCNKF_HUMAN | Potassium channel subfamily K member 15 |
| RRRXRSV | 2 | RRRSRSV | P18583 | SON_HUMAN | Protein SON |
| | | RRRKRSV | P49685 | GPR15_HUMAN | G-protein coupled receptor 15 |
| | | RRRASSV | O14681 | EI24_HUMAN | Etoposide-induced protein 2.4 homolog |
| RRRAXSV | 3 | RRRAQSV | Q7LDG7 | GRP2_HUMAN | RAS guanyl-releasing protein 2 |
| | | RRRAPSV | P21333 | FLNA_HUMAN | Filamin-A |
| | | RRRARPV | Q7RTU4 | BHA09_HUMAN | Class A basic helix-loop-helix protein 9 |
| RRRARXV | 4 | RRRARQV | Q8N9Z2 | CC71L_HUMAN | Coiled-coil domain-containing protein 71L |
| | | RRRARAV | Q6NUJ1 | SAPL1_HUMAN | Proactivator polypeptide-like 1 |
| | | RRRARVV | Q9GZQ6 | NPFF1_HUMAN | Neuropeptide FF receptor 1 |
| | | RRRARSW | Q8WUQ7 | CATIN_HUMAN | Cactin |
| RRRARSX | 6 | RRRARSS | P18825 | ADA2C_HUMAN | Alpha-2C adrenergic receptor |
| | | RRRARSP | Q96QZ7 | MAGI1_HUMAN | Membrane-associated guanylate kinase, WW and PDZ domain-containing protein 1 |
| | | RRRARSK | C9J069 | AJM1_HUMAN | Apical junction component 1 homolog |
| | | RRRARSR | O00198 | HRK_HUMAN | Activator of apoptosis harakiri |
| | | RRRARSL | Q9NZV5 | SELN_HUMAN | Selenoprotein N |
| TOTAL NUMBER | 19 | | | | |

| Motif | Number of proteins in UNIPROT contain the motif | MOTIF FOUND | PROTEIN Entry ID | PROTEIN Entry Name | PROTEIN full Name |
|---|---|---|---|---|---|
| PRRARSV | 0 | - | - | | |
| XRRARSV | 1 | ARRARSV | P37088 | SCNNA_HUMAN | Amiloride-sensitive sodium channel subunit alpha |
| PXRARSV | 0 | - | - | | |
| PRXARSV | 1 | PRPARSV | Q96PD5 | PGRP2_HUMA | N-acetylmuramoyl-L-alanine amidase |
| PRRXRSV | 1 | PRRSRSV | Q9UQ35 | SRRM2_HUMAN | Serine/arginine repetitive matrix protein 2 |
| PRRAXSV | 2 | PRRASSV | Q04844 | ACHE_HUMAN | Acetylcholine receptor subunit epsilon |
| | | PRRALSV | Q5VZ46 | K1614_HUMAN | Uncharacterized protein KIAA1614 |
| PRRARXV | 0 | - | - | | |
| PRRARSX | 1 | PRRARSS | Q92902 | HPS1_HUMAN | Hermansky-Pudlak syndrome 1 protein |
| TOTAL NUMBER | 6 | | | | |

| Motif | Number of proteins in UNIPROT contain the motif | MOTIF FOUND | PROTEIN Entry ID | PROTEIN Entry Name | PROTEIN full Name |
|---|---|---|---|---|---|
| SRRRARS | 0 | - | - | | |
| XRRRARS | 6 | RRRRARS | Q8WUQ7 | CATIN_HUMAN | Cactin |
| | | RRRRARS | P18825 | ADA2C_HUMAN | Alpha-2C adrenergic receptor |
| | | DRRRARS | Q96QZ7 | MAGI1_HUMAN | Membrane-associated guanylate kinase, WW and PDZ domain-containing protein 1 |
| | | PRRRARS | C9J069 | ALM1_HUMAN | Apical junction component 1 homolog |
| | | WRRRARS | O00198 | HRK_HUMAN | Activator of apoptosis harakiri |
| | | PRRRARS | Q9NZV5 | SELN_HUMAN | Selenoprotein N |
| SXRRARS | 3 | SDRRARS | Q8N2C7 | UNC80_HUMAN | Protein unc-80 homolog |
| | | SPRRARS | Q92902 | HPS1_HUMAN | Hermansky-Pudlak syndrome 1 protein |
| | | SCRRARS | Q5T4W7 | ARTN_HUMAN | Artemin |
| SRXRARS | 1 | SRDRARS | Q92917 | GPKOW_HUMAN | G-patch domain and KOW motifs-containing protein |
| SRRXARS | 1 | SRRQARS | Q9NSI2 | F2007A_HUMAN | Protein FAM207A |
| SRRRXRS | 10 | SRRRPRS | Q70EL4 | UBP43_HUMAN | Ubiquitin carboxyl-terminal hydrolase 43 |
| | | SRRRIRS | P05198 | IF2A_HUMAN | Eukaryotic translation initiation factor 2 subunit 1 |
| | | SRRRRRS | P18583 | SOV_HUMAN | Protein SON |
| | | SRRRSRS | Q8N2M8 | CLASR_HUMAN | CLK4-associating serine/arginine rich protein |
| | | SRRRSRS | Q15058 | KIF_HUMAN | Kinesin-like protein KIF14 |
| | | SRRRRRS | Q5M9Q1 | NKAPL_HUMAN | NKAP-like protein |
| | | SRRRSRS | Q14498 | RBM39_HUMAN | NA-binding protein 39 |
| | | SRRRSRS | Q96T37 | RBM15_HUMAN | RNA-binding protein 15 |
| | | SRRRSRS | Q13247 | SRSF6_HUMAN | Serine/arginine-rich splicing factor 6 |
| | | SRRRQRS | Q9UQ35 | SRRM2_HUMAN | Serine/arginine repetitive matrix protein 2 |
| SRRRAXS | 6 | SRRRAIS | Q00987 | MDM2_HUMAN | 3 ubiquitin-protein ligase Mdm2 |
| | | SRRRAQS | Q9NQU5 | PAK_HUMAN | Serine/threonine-protein kinase PAK 6 |
| | | SRRRADS | Q53GL0 | PKHO1_HUMAN | Pleckstrin homology domain-containing family O member 1 |
| | | SRRRAWS | Q9UKN7 | MYO15_HUMAN | Unconventional myosin-XV |
| | | SRRRAFS | Q9GZK7 | O11A1_HUMAN | Olfactory receptor 11A1 |
| | | SRRRAVS | Q9BYX2 | TBD2A_HUMAN | TBC1 domain family member 2A |
| SRRRARX | 3 | SRRRARR | O95450 | ATS2_HUMAN | A disintegrin and metalloproteinase with thrombospondin motifs 2 |
| | | SRRRARD | Q8N5L8 | RP25L_HUMAN | Ribonuclease P protein subunit p25-like protein |
| | | SRRRARV | Q9GZQ6 | NPFF1_HUMAN | Neuropeptide FF receptor 1 |
| TOTAL NUMBER | 30 | | | | |

| Motif | Number of proteins in UNIPROT contain the motif |
|---|---|
| XRRRARX | 47 |
| XRRRAXS | 44 |
| XRRRXRS | 139 |
| XRRXARS | 30 |
| XRXRARS | 22 |
| XXRRARS | 27 |
| SXRRARX | 29 |
| SXRRAXS | 29 |
| SXRRXRS | 72 |
| SXRXARS | 17 |
| SXXRARS | 20 |
| SRXRARX | 19 |
| SRXRAXS | 35 |
| SRXRXRS | 175 |
| SRXXARS | 16 |
| SRRXARX | 19 |
| SRRXAXS | 24 |
| SRRXXRS | 46 |
| SRRRXRX | 53 |
| SRRRXXS | 50 |
| SRRRAXX | 25 |
| TOTAL NUMBER | 938 |

| Motif | Number of proteins in UNIPROT contain the motif |
|---|---|
| RXXRS | 3774 |
| TOTAL NUMBER | 3774 |

The list of SLiMs of wild type and mutant C/H-CrUPs by the mutation P681R in SPIKE_SARS2 and detected in Human proteome are presented. Green block indicates the C/H-CrUP in wild-type protein, blue block indicates the mutant C/H-CrUP peptide by the P681R mutation and yellow block indicates the newly created C/H-CrUP by the same mutation. X (in red color) is used for the position within the peptide in order to create the motif. In the third column the detected motif is recorder, followed by the Protein Entry ID and the protein name in which it was detected. Total is summarizing the time in which the motifs related to the C/H-CrUP were recorded in the Human proteome.

**Suppl. Table 4. C/H-CrUPs of wild type and mutant Receptor-Binding Domain of SARS-Cov-2 Spike protein.**

| SARS-CoV-2 SPIKE PROTEIN RECEPTOR BINDING DOMAIN | | | | | | | | | | | | | | | |
|---|---|---|---|---|---|---|---|---|---|---|---|---|---|---|---|
| WILDTYPE | | | | | | | | MUTANT | | | | | | | |
| POSITION | C/H-CrUP | | | | | | Peptide number/ peptide length | MUTATION | VARIANT | NEW C/H-CrUP | | | | | Peptide number/ peptide length |
| 346 | VFNATR | FNATRF | NATRFA | ATRFAS | TRFASV | RFASVY | 6/6AA | R346K | Mu | VFNATK | FNATKF | ATKFAS | TKFASV | KFASVY | 5/6AA |
| 384 | YGVSPT | GVSPTK | VSPTKL | SPTKLN | PTKLND | | 5/6AA | P384L | Beta | YGVSLT | GVSLTK | SLTKLN | LTKLND | | 4/6AA |
| 417 | PGQTGKI | GQTGKIA | TGKIAD | GKIADY | | | 2/7AA, 2/6AA | K417N | Beta, Delta | GQTGNI | QTGNIA | TGNIAD | GNIADY | | 4/6AA |
| | | | | | | | | K417T | Gamma | PGQTGT | GQTGTI | QTGTIA | TGTIAD | | 4/6AA |
| 452 | GNYNYL | NYNYLY | NYLYRL | YLYRLF | LYRLFR | | 5/6AA | L452R | Alpha, Delta, Iota, Kappa | GNYNYR | YNYRY | NYRYRL | YRYRLF | | 1/5AA, 3/6AA |
| | | | | | | | | L452Q | Lambda | NYNYQ | YNYQY | NYQYRL | YQYRLF | QYRLFR | 2/5AA, 3/6AA |
| 478 | YQAGST | AGSTPC | STPCN | | | | 1/5AA, 2/6AA | T478K | Delta | YQAGSK | QAGSKP | AGSKPC | GSKPCN | KPCNG | 1/5AA, 4/6AA |
| 484 | CNGVEG | NGVEGF | GVEGFN | | | | 3/6AA | E484K | Alpha, Beta, Gamma, Eta, Mu | CNGVKG | NGVKGF | GVKGFN | KGFNC | | 1/5AA, 3/6AA |
| | | | | | | | | E484Q | Kappa | NGVQG | VQGFN | QGFNC | | | 3/5AA |
| 490 | FNCYF | CYFPLQ | YFPLQS | FPLQSY | | | 1/5AA, 3/6AA | F490S | Lambda | FNCYS | NCYSP | CYSPLQ | SPLQSY | | 2/5AA, 2/6AA |
| 494 | YFPLQS | FPLQSY | PLQSYG | QSYGF | SYGFQP | | 1/5AA, 4/6AA | S494P | Alpha | PLQP | LQPY | QPYG | PYGF | | 4/4AA |
| 501 | GFQPTN | FQPTNG | QPTNGV | PTNGVG | TNGVGY | NGVGYQ | 6/6AA | N501Y | Alpha, Beta, Gamma, Mu | GFQPTY | QPTYG | YGVGY | | | 2/5AA, 1/6AA |
| 516 | VVLSFE | VLSFELL | LSFELLH | FELLHA | ELLHAP | | 2/7AA, 3/6AA | E516Q | Beta | VVVLSFQ | VLSFQL | SFQLLH | FQLLHA | QLLHAP | 4/6AA, 1/7AA |

The C/H-CrUPs created by the mutations in the Receptor-Binding domain of SARS-CoV-2 wild-type and mutant spike protein (SPIKE_SARS2, P0DTC2) were identified. Peptide number/peptide length is the number of a given length C/H-CrUP around the position. By red color the amino acidss in wild-type C/H-CrUPs which will be replaced and the mutated amino acids in the new C/H-CrUPs were marked.

**Suppl. Table 5. C/H-CrUPs of mutant Receptor-Binding Motif of SARS-Cov-2 Spike protein contact positions.**

| CONTACT POSITION (#AA) | | | |
|---|---|---|---|
| WILDTYPE | MUTATION | VARIANT | NEW C/H-CrUP |
| T415 | | | |
| N439 | | | |
| Y449 | | | |
| Y453 | | | |
| F486 | | | |
| N487 | | | |
| Y489 | | | |
| Q493 | | | |
| Q498 | | | |
| T500 | | | |
| N501 | N501Y | Alpha, Beta, Gamma, Mu | GFQPTY QPTYG YGVGY |
| Y505 | | | |

The contact positions of Receptor-Binding Motif of wild type SARS-CoV- 2 spike protein with the ACE2 are shown in first column. Only the mutation N501Y was detected in these positions. The variants in which it appeared and the new C/H-CrUPs are listed. By red color the mutant amino acids are marked.

**Suppl. Table 6. NF9 C/H-CrUPS.**

| | PEPTIDES | | |
|---|---|---|---|
| | | MUTATION | |
| POSITION | SARS-CoV-2 | L452R | L452Q |
| 448 | NYNYLY | | NYNYQ |
| 449 | | YNYRY | YNYQY |
| 450 | NYLYRL | NYRYRL | NYQYRL |
| 451 | YLYRLF | YRYRLF | YQYRLF |

The C/H-CrUPs in wild-type and mutant NF9 peptide are listed. By red color the mutant amino acids were marked.
